# Pan-species transcriptomic analysis reveals a constitutive adaptation against oxidative stress for the highly virulent *Leptospira* species

**DOI:** 10.1101/2023.08.01.551416

**Authors:** Alexandre Giraud-Gatineau, Garima Ayachit, Cecilia Nieves, Kouessi C. Dagbo, Konogan Bourhy, Francisco Pulido, Samuel G. Huete, Nadia Benaroudj, Mathieu Picardeau, Frédéric J. Veyrier

**Author notes:** these authors contributed equally (Alexandre Giraud-Gatineau, Garima Ayachit). these authors contributed equally (Mathieu Picardeau, Frédéric J. Veyrier).

## Abstract

Different environments exert selective pressures on bacterial populations, favoring individuals with particular genetic traits that are well-suited for survival in those conditions. Evolutionary mechanisms such as natural selection have, therefore, shaped bacterial populations over time selecting, in a stepwise manner, the fittest bacteria that gave rise to the modern lineages that exist today. Advances in genomic sequencing, computational analysis, and experimental techniques continue to enhance our understanding of bacterial evolution and its implications. Nevertheless, these are often limited to genomic comparisons of closely related species. In the present study, we introduce Annotator-RNAtor, a graphical user interface (GUI) method for performing pan-species transcriptomic analysis and studying intragenus evolution. The pipeline uses third-party software to infer homologous genes in various species and highlight differences in the expression of the core-genes. To illustrate the methodology and demonstrate its usefulness, we focus on the emergence of the highly virulent *Leptospira* subclade known as P1+, which includes the causative agents of leptospirosis. Here, we expand on the genomic study through the comparison of transcriptomes between species from P1+ and their related P1-counterparts (low-virulent pathogens). In doing so, we shed light on differentially expressed pathways and focused on describing a specific example of adaptation based on a differential expression of PerRA-controlled genes. We showed that P1+ species exhibit higher expression of the *katE* gene, a well-known virulence determinant in pathogenic *Leptospira* species correlated with greater tolerance to peroxide. Switching PerRA alleles between P1+ and P1-species demonstrated that the lower repression of *katE* and greater tolerance to peroxide in P1+ species was solely controlled by PerRA and partly caused by a PerRA amino-acid permutation. Overall, these results demonstrate the strategic fit of the methodology and its ability to decipher adaptive transcriptomic changes, not observable by comparative genome analysis, that may have been crucial for the emergence of these pathogens.

**Author summary:** Natural selection is one of the central mechanisms of the bacterial evolution. Speciation events and adaptation occurs such as mutations, deletions and horizontal gene transfers to enhance our understanding of evolution. Nevertheless, these are often limited to genomic comparisons between species. Here, we are developed a graphical user interface method, named Annotator-RNAtor, to perform pan-species transcriptomic analysis and studying intragenus evolution. To illustrate the methodology, we focus on the emergence of the virulent *Leptospira* species, causative agents of leptospirosis. We shed light on a differential regulation of several PerRA-controlled genes in P1+ *Leptospira* subclade (highly virulent pathogens) compared to P1-*Leptospira* subclade (low virulent pathogens). P1+ species exhibit higher expression of the catalase-encoding gene *katE*, than P1-species, correlating with a greater ability to withstand peroxide. Additionally, we demonstrate that the difference in *katE* expression is mediated only by PerRA and the residue 89 of the PerRA protein participates on this regulation. These findings highlight the importance to decipher adaptative transcriptomic changes to fully understand the emergence of pathogenic species.

## Introduction

Bacteria are the most diverse and abundant cellular organisms on Earth, and in recent years, genomics has shed light on this diversity. The evolution of these species is influenced by their environment and the associated pressure of selection, driving natural selection. Events like mutations, deletions and horizontal gene transfers contribute to overall speciation events and adaptation [1]. One of the possible evolutionary trajectories includes establishing interactions with a mammalian host. It has been shown that the vast majority of host-associated symbionts evolved from free-living environmental counterpart [2]. The interplay between initially unrelated aspects of the ecology of microorganisms and a mammalian host can lead to a closer host–microbe symbiosis than can later evolve within the parasite-mutualist continuum [3, 4]. Understanding the genesis of the bacteria-host interaction is crucial for deciphering the causes of bacterial pathogenesis. This relationship requires, at its inception, the evolution of the bacterium to reach, survive and multiply at the site of infection. Next, the bacterium must acquire the mechanisms necessary to be maintained in the host population. This stepwise adaptation occurs through genetic alterations, with only permissive changes being selected at each of these steps. In this sense, several studies have described such ancestral evolution for several pathogens. For example, it has been suggested that current *Mycobacterium abscessus* clones circulating in humans have been first selected from an environmental pool of this species, followed by several rounds of within-host adaptation and transmission via the environment. This stepwise evolution could have been mediated by horizontal transfer of genes and more specifically genes encoding global transcriptional regulator that mediated initial phenotypic variance [5].

With advances in NGS sequencing technologies, it has become easier to infer and mine information from genomic data. Nevertheless, studies have also highlighted the importance of transcriptomic comparison in discerning the mechanisms of regulatory evolution in several bacteria [5-7]. These studies indicate the importance of transcriptomics, but they are limited to comparisons between closely related bacteria. Despite the widespread use of RNAseq as a method to study gene expression changes and the existence of several tools for investigating differential expression of genes between two groups, there are limited available tools or pipelines for studying differential expression in multiple species simultaneously. It is therefore challenging to compare more diverse bacteria to investigate ancestral events, such as those linked to the emergence of clades or subclades of bacteria within a genus (herein referred to as intragenus evolution). Nevertheless, a recent study has successfully compared the expression of genes between bacteria of the same family that differ in their cell shape. This analysis has revealed the effect of the loss of the division and cell wall (*dcw*) cluster regulator MraZ in multicellular and longitudinally dividing *Neisseriaceae* MuLDi [8]. Similarly, another study reported the ancestral nitrogen-fixing root nodule symbionts transcriptome by combining transcriptomics and phylogenomics of multiple species [9].

In the present study, we introduce Annotator-RNAtor, a graphical user interface (GUI) method for studying intragenus transcriptomic evolution. The pipeline can infer homologous genes in various species and highlight differences in the expression of the core-genes. To illustrate the methodology and demonstrate its usefulness in deciphering intra-genus evolution, we focus on the emergence of the highly virulent *Leptospira* subclade, known as P1+ (sometimes referred to as P1hv for highly virulent pathogens in animal models). This subclade includes the multiple causative agents of leptospirosis, an emerging zoonotic disease transmitted to humans through exposure to soil or water contaminated with the urine of animal reservoirs. Annually, an estimated 1 million cases of leptospirosis and nearly 60,000 deaths occur, resulting in a loss of 2.9 million disability-adjusted life years [10, 11]. These numbers are likely to be underestimations due to misdiagnosis and under-reporting, particularly in regions where other diseases with similar non-specific presentations, such as dengue and malaria, are prevalent [12-16]. Leptospirosis is becoming even more prevalent due to global climate changes that cause more frequent and severe flooding events. Notably, leptospirosis is not restricted to tropical regions and developed countries are also experiencing the occurrence of the disease in human and animals [17, 18]. Recent elaborated phylogenetic analyses have resulted in a comprehensive new genome-based classification scheme of the *Leptospira* genus [19]. Interestingly, this study described the genomic features of the specific sublineage of *Leptospira* P1+, which diverged after a specific node of evolution and that is most often associated with severe infections in humans and animals (node 1 in Fig 1) [19, 20]. Here, we expand on the genomic study by investigating the cause of the emergence of this lineage through the comparison of transcriptomes between species from the P1+ lineage and their related P1-counterparts (or P1lv for low-virulent pathogens in animal models, see Fig 1). In doing so, we shed light on several pathways differentially expressed and specific disparity in the regulation of several PerRA-regulated genes in P1+ species. PerRA is a transcriptional regulator of the Fur family that regulates the adaptation to oxidative stress in *Leptospira* [21, 22]. PerRA represses the catalase-encoding gene *katE* and genes encoding the peroxidase AhpC and the cytochrome C peroxidase CCP. These enzymes have been identified as the first line of defense against peroxide in *L. interrogans* [22]. The PerRA regulon also contains genes whose expression is activated by this regulator, including a cluster encoding the TonB-dependent energy transduction system [22, 23]. Here, we show that P1+ species exhibit higher expression of the *katE* gene, a well-known leptospiral virulence determinant [24], correlating with a greater catalase activity and ability to withstand peroxide. Our findings indicate that the higher *katE* expression in P1+ species is due to a basal *katE* semi-derepression caused by amino-acid permutations of PerRA. Overall, these results demonstrate the strategic fit of the methodology, in parallel of intragenus genomic comparisons, and its ability to decipher transcriptomic changes that may have been crucial for the emergence of these pathogens.

**Fig 1.**
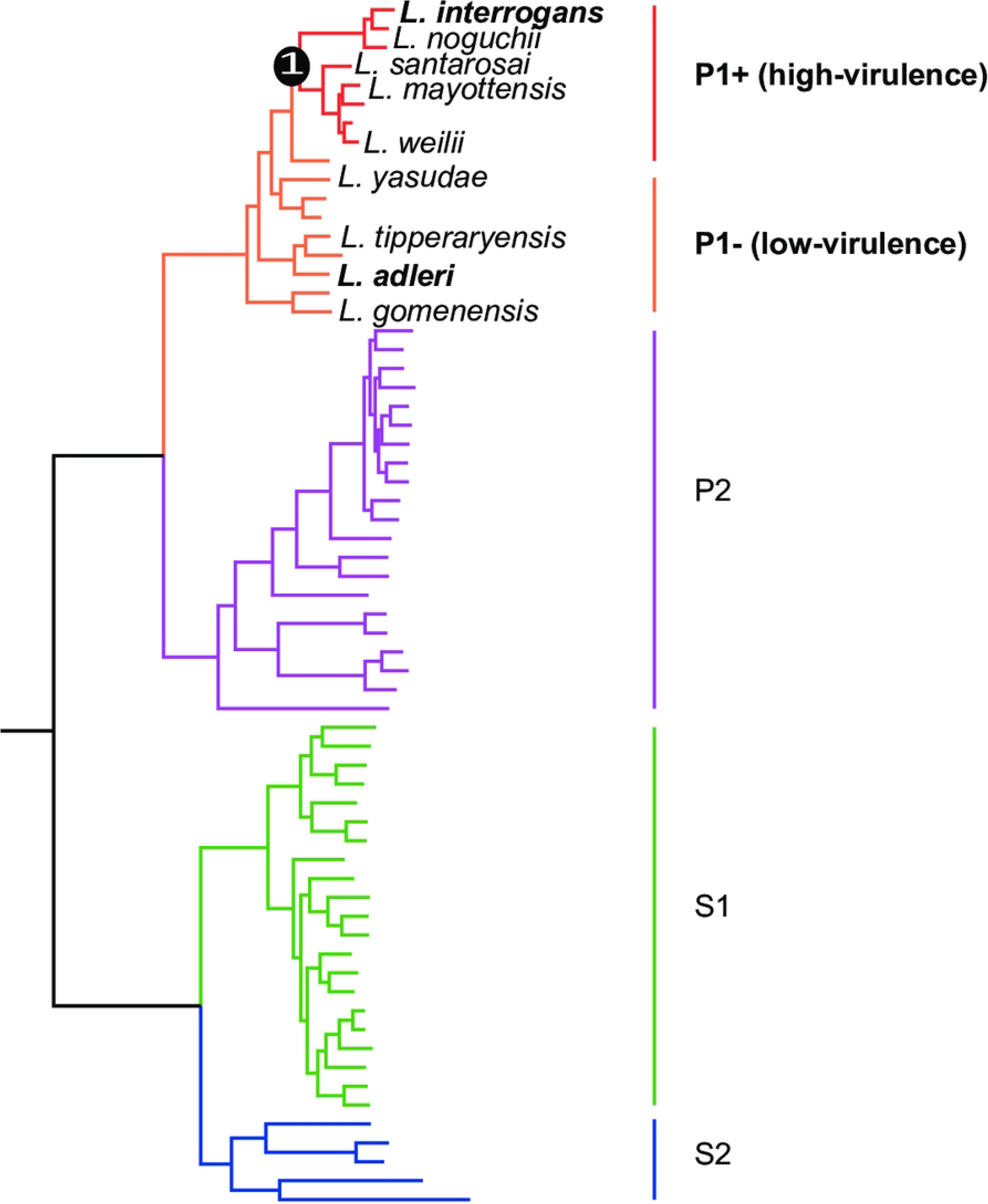
The subclade P1, formerly referred to as the “pathogens” lineage, can be separated into two distinct groups: P1+ and P1-. P1+ consists of species associated with severe infections and diverged after a specific node of evolution (node 1), while P1-comprises species that have not been isolated from patients and are considered as “low-virulent pathogens”.

## Methods

### Bacterial strains and culture conditions

The *Leptospira* strains used in this study (as listed in S1 Table) were cultivated aerobically in Ellinghausen-McCullough-Johnson-Harris medium (EMJH) at 30°C with shaking at 100 rpm. The *E. coli* strains β2163 and Π1 were cultivated at 37°C with shaking, in Luria-Bertani medium supplemented with 0.3 mM thymidine or diaminopimelic acid, respectively.

### RNA sequencing and comparisons

To assess the performance of the pipeline, a comparative analysis was conducted on nine genomes of the genus *Leptospira* (Fig 1 and Table 1). Both genomic and transcriptomic data for these strains were generated in-house. For all experiments, *Leptospira* species were used during the exponential phase of growth. *Leptospira* strains on exponential phase cultures were resuspended in TRIzol lysis reagent (ThermoFisher Scientific). Nucleic Acids were extracted with chloroform and precipitated with isopropanol as previously described [25] and DNA was removed by DNAse treatment using the Turbo DNA-free kit (Thermo Scientific^TM^). Prior to cDNA synthesis (RevertAid RT Reverse Transcription Kit, K1691, Thermo Scientific^TM^), rRNA was removed from 1 μg of total RNA using the NEBNext rRNA Depletion Kit. The sequencing was performed at the Genome Québec Innovation Centre (McGill University, Montréal, Canada). Briefly, sequencing libraries were constructed using the Illumina® Stranded mRNA Prep Kit according to the manufacturer’s instructions. Subsequently, 100 bp pair-end sequencing was carried out using the NovaSeq 6000 System. The FastQ reads have been deposited in the SRA database PRJNA998607. Sequence reads were initially processed with FastQC (version 0.73) to assess their quality before being trimmed using FastQ Groomer (version 1.1.5).

**Table 1.**
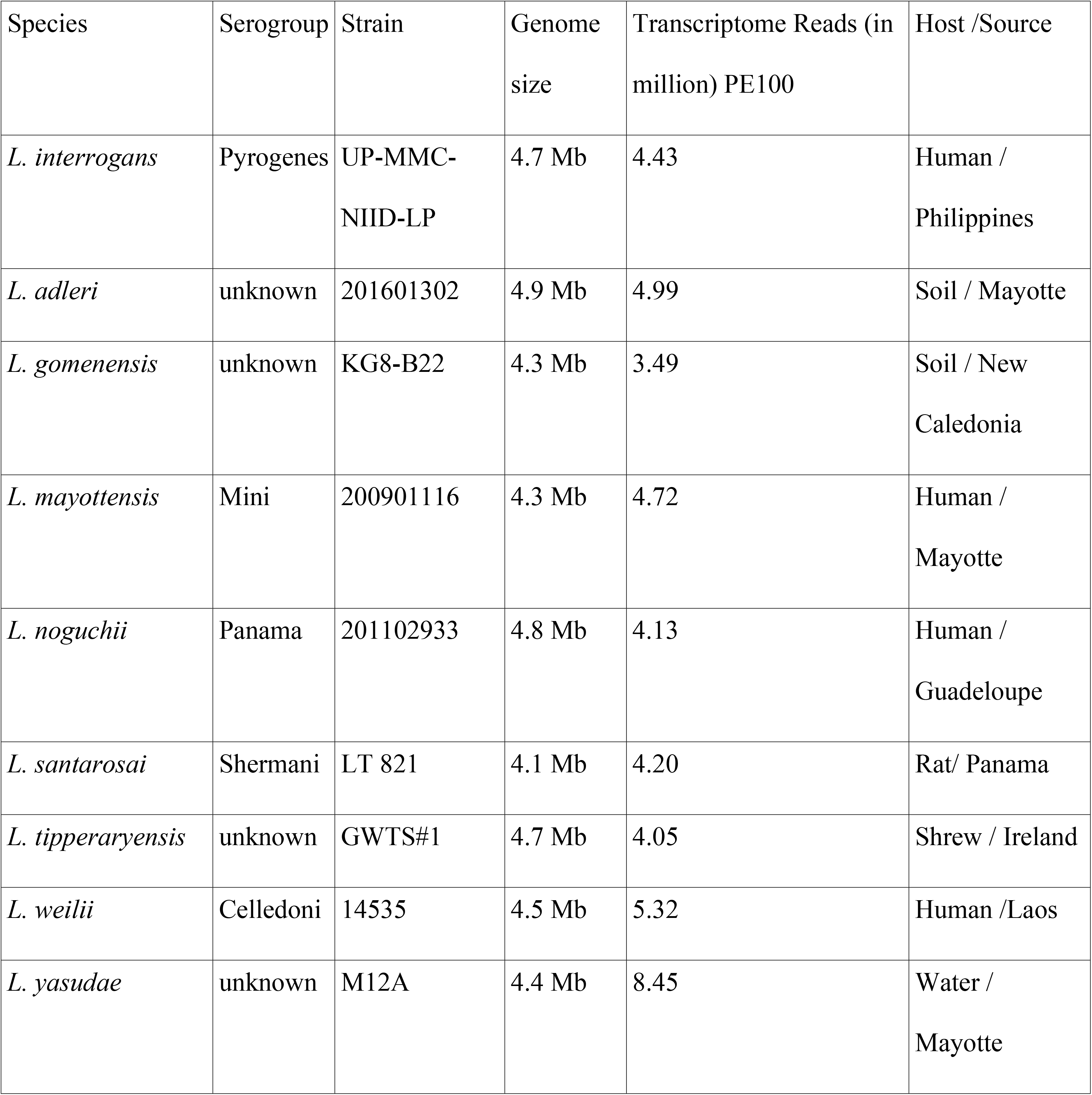
Details of species used for testing Annotator-RNAtor.

| Species | Serogroup | Strain | Genome size | Transcriptome Reads (in million) PE100 | Host /Source |
| --- | --- | --- | --- | --- | --- |
| *L. interrogans* | Pyrogenes | UP-MMC-NIID-LP | 4.7 Mb | 4.43 | Human / Philippines |
| *L. adleri* | unknown | 201601302 | 4.9 Mb | 4.99 | Soil / Mayotte |
| *L. gomenensis* | unknown | KG8-B22 | 4.3 Mb | 3.49 | Soil / New Caledonia |
| *L. mayottensis* | Mini | 200901116 | 4.3 Mb | 4.72 | Human / Mayotte |
| *L. noguchii* | Panama | 201102933 | 4.8 Mb | 4.13 | Human / Guadeloupe |
| *L. santarosai* | Shermani | LT 821 | 4.1 Mb | 4.20 | Rat/ Panama |
| *L. tipperaryensis* | unknown | GWTS#1 | 4.7 Mb | 4.05 | Shrew / Ireland |
| *L. weilii* | Celledoni | 14535 | 4.5 Mb | 5.32 | Human /Laos |
| *L. yasudae* | unknown | M12A | 4.4 Mb | 8.45 | Water / Mayotte |

**Table 2.** Homologous genes (core genome) identified and reannotated in all P1 *Leptospira* genomes.

|  |  |
| --- | --- |
| NetworkX | GET_HOMOLOGUES |
| 2370 | 2323 |

### Software Annotator-RNAtor for RNA comparisons

#### *Overall analyses* (Fig 2)

Further comparative expression analyses were conducted as follows. The genome sequences of the nine evaluated *Leptospira* species (Table 1), were annotated using Prokka v1.14.5. For each dyad possibility, a standalone BLASTP search was performed using the respective protein sequences (.faa) files. Network connections were established using a python programming package NetworkX version 2.6.2[26], with a similarity threshold set at 60%. Essentially, all proteins showing more than 60% similarity with one of the members (putative homologues) were clustered together. Each cluster of proteins was assigned a name (e.g., Lsp_1), which was then used to replace the original locus tags in the .gff file generated by Prokka. Protein files were also screened for putative homologues using GET_HOMOLOGUES [27] through OrthoMCL method. The reannotated .gtf files were utilized to map the reads to their corresponding genomes using BWA.

**Fig 2.**
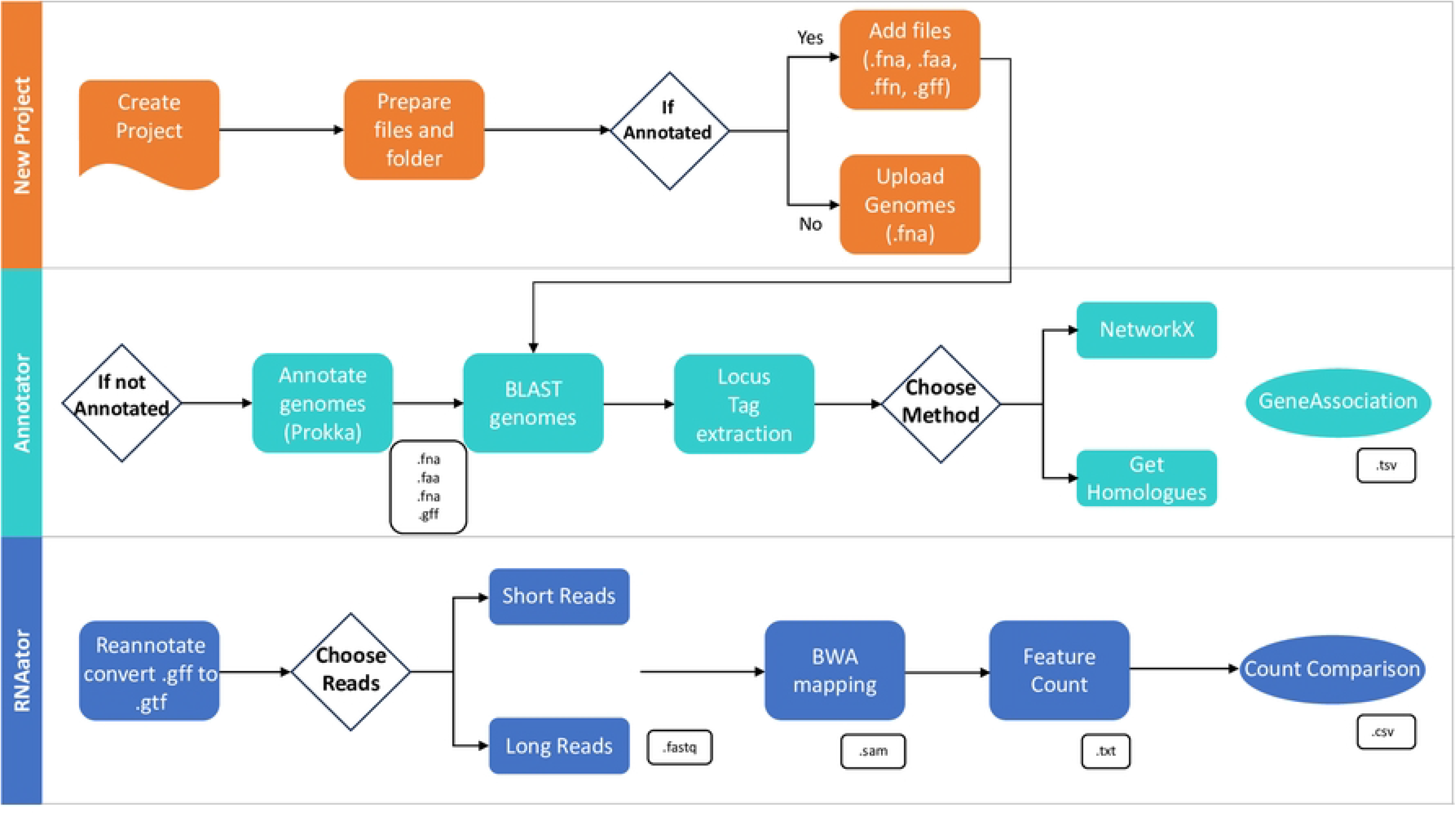
Pipeline workflow and modules.

#### Input Data and formats using the GUI

The pipeline accepts whole-genome sequence FASTA files. If the genomes are already annotated, it requires genome FASTA, protein FASTA, genome feature files and coding sequences for Annotator. The RNAtor module requires related FASTQ files as input. Currently, the pipeline supports analysis of both paired-end short-read as well as long-read nanopore data.

#### Third party software and dependencies, data processing and clustering

The pipeline utilizes Prokka for annotating the genomes, followed by BLAST+ programs [28, 29]. NetworkX [26] and GET_HOMOLOGUES v16052022 [27] are used for screening homologues in subsequent steps [27]. Homologues can be screened based on the percent similarity/identity as parameters. Additionally, GET_HOMOLOGUES provides options to cluster homologues using Bidirectional best-hit, COGtriangles v2.1[30], or OrthoMCL v1.4 [31]. At the end of the process, a tab-separated file is generated, in which Annotator-RNAtor creates a presence (locus) or absence (-) map, facilitating the finalization of the analyses. The RNAtor menu in Annotator-RNAtor implements the bwa [32] and bwa-mem commands. Annotator-RNAtor is designed to utilize the annotation files from NCBI. RNAtor invokes Pandas and FeatureCounts of Subread [33].

#### Supported platforms and availability

Annotator-RNAtor is built using Python3 with PyQt bindings for a user-friendly GUI interface. The efficiency of Annotator-RNAtor relies on running tasks in parallel, utilizing multiple processors and GNU Parallel. The pipeline has been tested on Linux/Ubuntu with 8 GB RAM and 64-bit processors. It includes a user manual with a detailed hands-on tutorial and an installation script that verifies dependencies. Alternatively, due to constant version upgrades in software, the package also includes a conda recipe file to install required dependencies in conda environment. The stand-alone pipeline can be downloaded from GITHUB via https://github.com/BactSymEvol/Annotator.

#### Statistical analyses

In the RNAtor module, the .gtf and .sam files were utilized to perform read counts using featureCounts v2.0.1 from the Subread package. The count files for each sample were combined into a table using a custom script, and these results were subsequently analyzed using DESeq2 version 3.14 [34]. To retain only genes from the soft-core genome and exclude genes from shell and cloud genomes, genes without any counts in less than 8 species were not considered. The genomes were grouped into P1+ and P1-categories. The data were normalized and transformed using the VarianceStabilizingTransformation (VST) function to yield a homoscedastic matrix of values. Further, Log_2_ fold change values were extracted with an FDR-adjusted p values (padj) cut-off of 0.05. To facilitate the interpretation of results, the differentially regulated genes are presented using the locus tag of *Leptospira interrogans* serovar Manilae str UP-MMC-NIID-LP. We investigated the biological functions of the gene differentially expressed (Fold change>2 and padj <0.05) and the putative pathways that could link them through a STRING [35] association analysis and COG [36]. The STRING analysis enabled the identification of protein interactions and functional enrichment in the two datasets, which consisted of up- and down-regulated genes in P1^+^ species. To accomplish this, the protein-coding sequences from both datasets were concatenated and analyzed collectively using STRING v11.5 [35], with *L. interrogans* serovar Lai str 56601 serving as the reference. Of note, *L. interrogans* serovar Manilae, which had been the reference genome throughout the study, is not yet included in the STRING database. For the analysis, the highest confidence interaction score of 0.900 was employed to focus solely on strong interactions, and any disconnected nodes (protein) in the network were hid. On the other hand, the protein-coding sequences from both datasets were classified based on Clusters of Orthologous Genes (COG) using eggNOG mapper (options *--evalue 0.001 --score 60 --pident 40 --query_cover 20 --subject_cover 20 -- target_orthologs all --pfam_realign denovo*) [37, 38]. The representativeness of COG categories in each dataset was calculated by determining the ratio of protein-coding genes in each category, normalized by the total number of protein-coding genes in *L. interrogans* serovar Manilae (3,572).

### Quantitative reverse transcription PCR (RT-qPCR)

Total RNAs from *Leptospira* strains were extracted using QIAzol lysis reagent (Qiagen) and purified with RNeasy columns (Qiagen). Reverse transcription of mRNA to cDNA was carried out using the iScript™ cDNA Synthesis kit (Bio-Rad), followed by cDNA amplification using the SsoFast™ EvaGreen® Supermix (Bio-Rad). All primers used in this study are listed in S2 Table. Reactions were performed using the CFX96 real-time PCR detection system (Bio-Rad). The relative gene expression levels were assessed according to the 2^-ΔCt^ method using *flaB2* (LIMLP_09410) as reference gene.

### Determination of bacteria viability and catalase activity

2.10^8^ of exponentially growing *Leptospira* species and *L. interrogans* Δ*katE* were incubated in EMJH with or without 1 mM of H_2_O_2_ for 30 min at 30°C. Rezasurin (Alamar Blue Assay, ThermoFisher) was added, and bacteria were further incubated for 24h. The absorbance was measured at 540 nm. Viability was determined based on the ability of cells to reduce rezasurin into resorufin. The percentage of cell viability was calculated as the ratio of rezasurin reduction for bacteria incubated with H_2_O_2_ to bacteria incubated in the absence of H_2_O_2_ as described previously [39]. For colony-forming unit (CFU) determination, *Leptospira* (treated and untreated with H_2_O_2_) were diluted in EMJH and plated on EMJH agar plates. Colonies were counted and the percent survival (% of CFU) was calculated as the ratio of CFU for bacteria incubated with H_2_O_2_ to bacteria incubated without H_2_O_2_. A bacterial culture containing about 10^9^ exponentially growing *Leptospira* was used to determine the catalase activity using the Catalase Activity Assay Kit (Abcam), according to manufacturer’s instructions.

### PerRA alignment analysis

Ortholog sequences of the PerRA protein in *Leptospira species* were searched with BLASTP version 2.13.0 and HMMer version 3.3.1 against the reference proteomes database of the 68 *Leptospira* spp. Only the sequences with an e-value ≤ 0.01 were retained. The results were subsequently aligned by MAFFT version 7.467 using the *L-INS-I* algorithm, and tree inference was computed with IQ-TREE version 2.0.6 with the best-fit model JTTDCMut + F + R5. Direct homologues of PerRA were extracted from this phylogeny, and the final alignment was refined using MAFFT (version 7.467 using the *L-INS-i* algorithm) with only these sequences. The alignment of PerRA among *Leptospira* spp. was visualized with Jalview software (using Clustal for sequence coloring [40]). Multiple sequence alignments of all P1+ and P1-species were visualized using *Alvis* software version 0.1. The residues of PerRA involved in the coordination of the regulatory metal were represented using Mol* 3D Viewer of PDB website.

### Inactivation of *perRA* of *L. adleri*

*L. interrogans* Δ*perRA* was obtained as previously described [22]. Inactivation of *perRA* gene in *L. adleri* was performed by allelic exchange, replacing the *perRA* coding sequence with a kanamycin resistance cassette. The kanamycin resistance cassette flanked with 0.8 kb sequences homologous to the adjacent *perRA* sequences was obtained by gene synthesis (GeneArt, Life Technologies), and subsequently cloned into a kanamycin-resistant *E. coli* vector. The resulting suicide plasmid (pSΔ*perRA*, S3 Table) was introduced into *L. adleri* by electroporation, as previously described [41], by Gene Pulser Xcell (Bio-Rad) and transformants were plated on EMJH supplemented with 50 µg/mL kanamycin. Individual kanamycin-resistant colonies were selected and screened by PCR (using PERADL1and PERADL2 primer set, listed in S2 Table).

### Complementation of *Leptospira perRA* mutants

*PerRA* expression in all *perRA* mutants were performed with the leptospiral replicative vector pMaOri [42] (S3 Table). A vector containing the *L. interrogans perRA* gene under the control of its native promoter (pNB138 named here *perRA*_P1+_) was previously obtained [42, 43]. The vector harboring the *L. adleri perRA* gene, along with its native promoter region (200 bp upstream region), was amplified from genomic DNA of *L. adleri* (using ComAdPerRA1 and ComAdPerRA2 primer set, S2 Table) and cloned between the SacI and XbaI restriction sites in the pMaORI vector. The absence of mutations in the *perRA* locus in the resulting plasmid (perRA_P1-_, S3 Table) was confirmed by DNA sequencing (Eurofins). Then, the *perRA*_P1-_ plasmid was introduced into *L. interrogans* Δ*perRA* and *L. adleri* Δ*perRA,* and the *perRA*_P1+_ plasmid was introduced into the *L. adleri* Δ*perRA* by conjugation using the *E. coli* β2163 conjugating strain, as previously described [44]. *Leptospira* conjugants were selected on EMJH agar plates containing 50 µg/mL spectinomycin.

### Mutagenesis of *perRA* sequence

The H89N substitution of *L. interrogans* perRA ORFs in *perRA*_P1+_ was conducted using the QuikChange Lightning Multi Site-Directed Mutagenesis Kit (Agilent) according to the manufacturer’s protocol and the mutagenic oligonucleotide primer PerRA_c265a. (S2 Table). The presence of the c265a mutation and the absence of additional mutation in the *perRA* locus in the obtained plasmid (*perRA*_P1+-H89N_) was confirmed by DNA sequencing (Eurofins). Subsequently, the plasmid *perRA*_P1+-H89N_ was introduced into *L. interrogans* and *L. adleri* Δ*perRA* mutants by conjugation using the *E. coli* β2163 strain.

### Western Blot analysis

Total extracts of *Leptospira* were obtained by sonication in a lysis buffer containing 25 mM Tris pH 7.5, 100 mM KCl, 2 mM EDTA, 5 mM DTT, and a protease inhibitors cocktail (cOmplete Mini EDTA-free, Roche). A total of 7.5 µg of each lysate were resolved on a 15% SDS-PAGE and transferred onto nitrocellulose membrane. PerRA was detected by immunoblot, as described previously[43].

### *In vivo* animal studies

Protocols for animal experiments conformed to the guidelines of the Animal Care and Use Committees of the Institut Pasteur (Comité d’Ethique d’Expérimentation Animale CETEA # 2016–0019), agreed by the French Ministry of Agriculture. All animal procedures carried out in this study complied with the European Union legislation for the protection of animals used for scientific purposes (Directive 2010/63/EU). Male 4 weeks-old Syrian Golden hamsters (RjHan:AURA, Janvier Labs) were infected (4 per group) by intraperitoneally injection with bacterial suspensions containing 10^6^ or 10^8^ for *L. interrogans* or *L. adleri*, respectively, as enumerated using a Petroff-Hausser counting chamber. The animals were monitored daily and euthanized by carbon dioxide inhalation upon reaching the predefined endpoint criteria (sign of distress, morbidity). To assess leptospiral load, blood, kidney, and liver were sampled and DNA was extracted with the Tissue or Blood DNA purification kit (Maxwell, Promega). The burden in blood and tissues was determined by qPCR with the Sso Fast EvaGreen Supermix assay (Bio-Rad) using the *flaB2* gene, and the concentration of host DNA was quantified using the *gapdh* gene. *Leptospira* load was expressed as genomic equivalent (GEq) per µg of host DNA.

### Quantification and statistical analysis

Data are expressed as means ± standard deviations (SD), of at least 3 independent biological replicates. Statistical analyses were performed with Prism software (GraphPad Software Inc.), using the *t* test as indicated in the figure legends.

## Results

### General description of Annotator and RNAtor workflow

There are two primary modules to analyse data in this pipeline (Fig 2). In the initial steps, Annotator module assigns homologous sequences in the different species studied and reannotates the genomes. Annotator uses two different methods namely NetworkX [26] and GET_HOMOLOGUES [27] to assign homology. The user can define similarity cut-off for NetworkX and identity cut-off for GET_HOMOLOGUES. This will depend on the phylogenetic proximity of the strains studied (herein we use 60% and 45% respectively). In the end, the pipeline reannotates the homologous genes with common user-defined unique tags that are found in the different species (convert GFF to GTF). A gene association table is generated at the end of this module which shows presence-absence of genes in the genomes (S4 Table). Singletons are not reannotated using the pipeline and can be seen as locus tags in the table. Once the gene association table is generated, the RNAtor module is used to map transcriptome data onto the genomes to generate a counts matrix file. This matrix is generated using the .GTF files (containing reannotated common unique names), BWA for mapping and featureCounts scripts to count the number of reads by genes. The final counts comparison file (S4 Table) shows the expression of the homologous genes in all the species studied. Species-specific gene counts can also be seen.

### Inferring homology using NetworkX or GET_HOMOLOGUES

Multiple tools exist to perform the analysis all of which all have their advantages and disadvantages. For example GenAPI [45], PanSeq [46], Roary [47], SaturnV [48], PanDelos [49] and panX [50] to name a few. Some tools are based on alignment of the query genome sequence to a reference genome to specifically test for the presence-absence of genes that are present in the given reference. Other tools construct a pan-genome from genome assemblies and then determine the gene set present in each of the genome assemblies. The performance of this approach depends on the completeness of the queried genome sequences, and all previously mentioned tools are designed for analysis of near-complete genomes with only minor parts of genes missing due to sequencing and assembly imperfections. Apart from these, Hidden Markov approach also exist which can be used to detect homologies but may require higher computational power and well-curated datasets. In the current pipeline, homology analysis is performed using the Annotator module which considers protein similarity searches instead of DNA, making it more sensitive. DNA:DNA alignments cannot detect homology post divergence of 200-400 million years. Compared to this, protein:protein or protein:translated alignment are more accurate. Protein:protein alignments with expectation values < 0.001 can reliably be used to infer homology, whereas DNA:DNA expectation values < 10^−6^ often occur by chance, and 10^−10^ is a more widely accepted threshold for homology based on DNA:DNA searches [51].

We used two methods relying on BLAST to search for homologues. NetworkX [26] uses a graph method to screen out homologues from the BLAST results (based on the user-provided threshold) whereas GET_HOMOLOGUES [27] uses identity to screen them. These approaches both have their benefits depending on the type of data being analysed. For example, if the purpose of the analysis is to find gene presence/absence in unrelated species or genera the NetworkX method provides more flexibility when searching for similar sequences. It also takes into account fragmented proteins that are not considered as homologues by other tools. This may provide an advantage while working with fragmented assemblies as well. In this study, we analyzed 9 *Leptospira* genomes (Fig 1 and Table 1) along with their transcriptome data. Annotator module was used to extract homologous genes and generate a gene presence/absence table using both NetworkX and OrthoMCL in GET_HOMOLOGUES method. In total we obtained 2370 core genes using NetworkX (60% similarity) and 2323 core genes using GET_HOMOLOGUES (45% identity). The difference comes from the fact that the NetworkX method is based on similarity between sequences and considers fragmented sequences as well whereas GET_HOMOLOGUES is based on identity and has a more stringent approach. GET_HOMOLOGUES does not take into account fragmented or dissimilar sequences (such as those found when comparing divergent species) and annotates them separately [27]

### Transcriptomic comparison between *Leptospira* species using RNAtor

The reannotated genomes by the two methods generated two new .GTF files. These files were used to map the reads to their corresponding genomes using BWA and the number of reads per gene was determined using featureCounts, which produced a .csv table containing the reannotated genes and their respective counts (S4 Table). For inferring differential expression between the P1+ and P1-subclades, standard DESeq2 analysis was employed, excluding genes that were absent or not expressed in more than one species. A total of 2323 and 2370 instances were analyzed using data from GET_HOMOLOGUES and NetworkX, respectively (S4 Table). The volcano plot (Fold change in function of the padj values) in Figure 3 presents the results obtained with NetworkX (Fig 3A) and GET_HOMOLOGUES (Fig 3B). Notably, both methods yielded highly similar results, as the majority of over-expressed and under-expressed genes in the P1+ subclade were identified by both methods (see Venn Diagram Fig 3D). NetworkX detected approximately 10% more genes overall. The slight variations in the expression values between NetworkX and GET_HOMOLOGUES may be attributed to the separate reannotation processes and the differences in the parameters utilized (% similarity vs identity, respectively). Fig 3C presents a circular representation of results obtained for Chromosome 1. The central inner circles represent genes that are not distributed in all the species (cloud or shell genomes), thus excluded from the analyses. Of note, this representation highlights some non-conserved regions with content variation between species (such as exemplified with the *rfb* cluster of genes in Fig 3C). The outer circles depict fold change values obtained with both methods, revealing only subtle differences between the two approaches.

**Fig 3.**
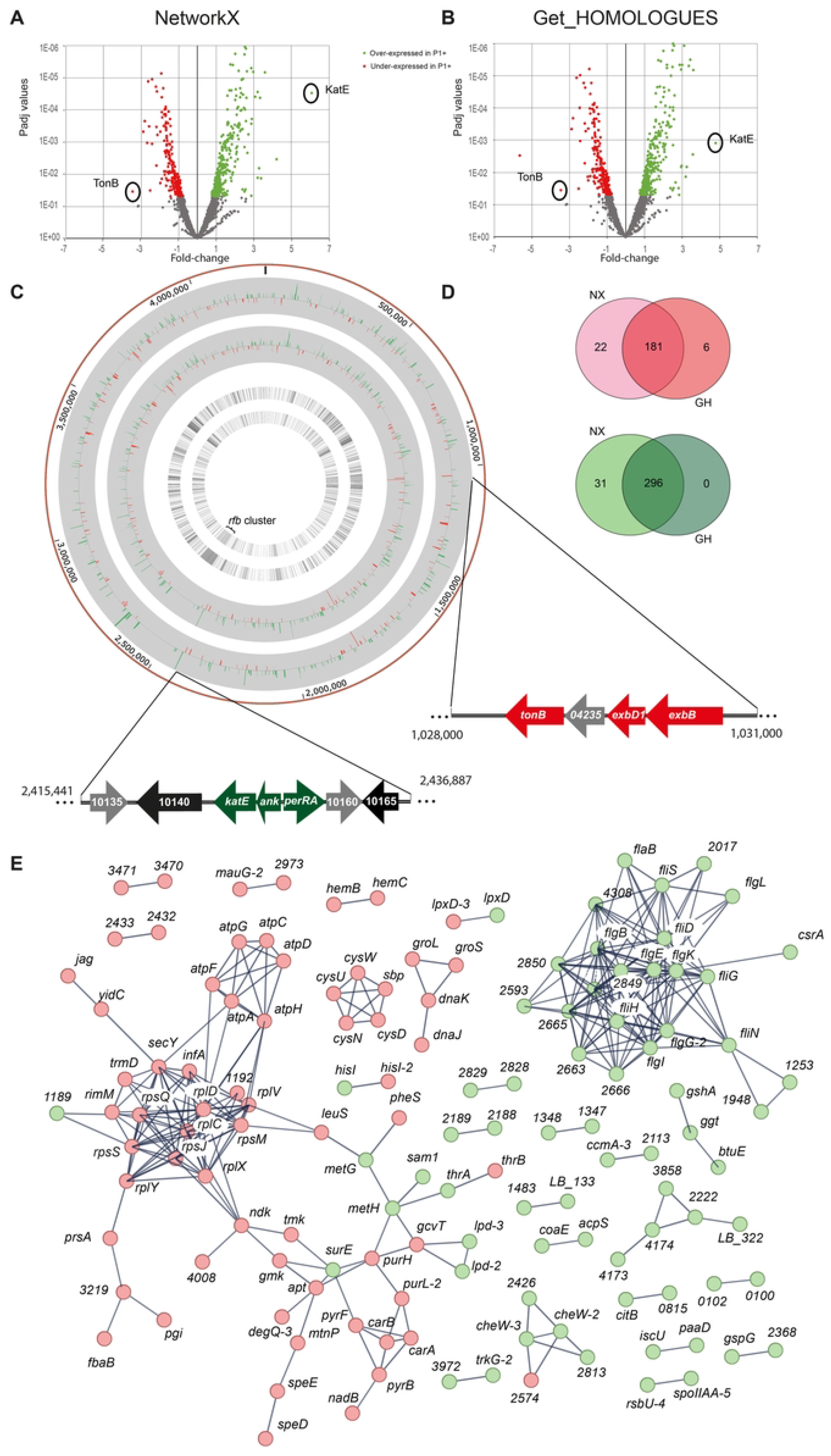
Comparison of differentially expressed genes in P1+ species compared to P1-using both methods. Volcano plot of RNAseq analysis of P1+ species compared to P1-. Using NetworkX in (**A**) and GET_HOMOLOGUES in (**B**). *p* value is plotted against fold change and were calculated using DeSeq2. Red points represent genes under-expressed and green genes over-expressed in P1+ as compared to P1-. (**C**) Circos plot generated for chromosome 1. The outermost track shows the chromosome, the second shows gene expression values generated using GET_HOMOLOGUES followed by expression values generated using NetworkX. The second innermost track show genes with no homologues in other species determined using GET_HOMOLOGUES and the innermost track using NetworkX. Finally, the *perRA* and *tonB* loci are also indicated with color coded genes (grey: no homologues in other species, black: Log2FC between 1.5 and -1.5, red: Log2FC<-1.5, green: Log2FC>1.5) (**D)** Venn diagram showing the comparison of the number of genes over-expressed (red) or under-expressed (green) in P1+ obtained with both methods (NX for NetworkX and GH for GET_HOMOLOGUES). **(E)** STRING association analysis. In red are transcripts that are less abundant in P1+ and in green are transcripts that are more abundant in P1+ as compared to P1-. Locus_tag from *Leptospira interrogans* serovar Lai str. 56601 are used as reference by STRING (such as 1483 for LA_1483).

Concerning the differentially expressed genes identified using this methodology, we used a STRING association analysis to visually shed light on pathways that could be differentially expressed between two groups (Fig 3E). From this, a clear overexpression of genes encoding the flagellation machinery was observed in P1+ and an underexpression of pathways from the general metabolisms. Indeed, the functional COG [36] analysis revealed a significantly higher representation (20-fold) of the cell motility category in the overexpressed genes compared to the underexpressed ones (S5 Table). There is also a 3.5-fold enrichment observed in the category of signal transduction mechanisms among the overexpressed genes in comparison to the underexpressed. Conversely, categories involved in the transport and metabolism of amino acids, nucleotides, carbohydrates, lipids, and coenzymes are more represented in the underexpressed genes, indicating potential affected metabolic pathways in the P1+ species.

To validate the method, our focus was solely on genes that exhibited the most differential expression between the P1+ and P1-groups. Interestingly, we observed two genes, *tonB* and *katE*, which are part of two distant loci, showing altered expression between the two groups (see *ank/perR* and *exbD1/exbB* genes respectively in Fig 3C). Intriguingly, both loci are part of the PerRA regulon, with *tonB* and *katE* being respectively the most underexpressed and overexpressed genes in a *L. interrogans perRA* deletion mutant [21, 22]. Importantly, most of the genes detected differentially expressed in this mutant were also differentially expressed between P1+ and P1-(S6 Table). Most of the genes from this regulon are known to be modulated in the presence of peroxide to protect the cell against oxidative stress. Therefore, our results strongly suggest a disparity in the expression of the PerRA regulon between species from the P1+ and P1-subclades.

### A differential expression of PerRA-controlled genes is associated with an increased resistance to peroxide in P1+

The differential expression of the PerRA-controlled genes (some positively, others negatively) detected with RNAtor between P1+ vs P1- was confirmed by RT-qPCR. Consistent with our transcriptomic data, we observed an increase in the expression of the *ank*-*katE* operon in P1+ species compared to P1-species (20.2-fold and 47.5-fold increase, respectively, Fig 4A). Similarly, we confirmed by RT-qPCR that the *tonB* locus (*exbD1*, *exbD2* and *tonB*), which is known to be positively regulated by PerRA [21, 22], was under-expressed in P1+ (Fig 4B).

**Fig 4.**
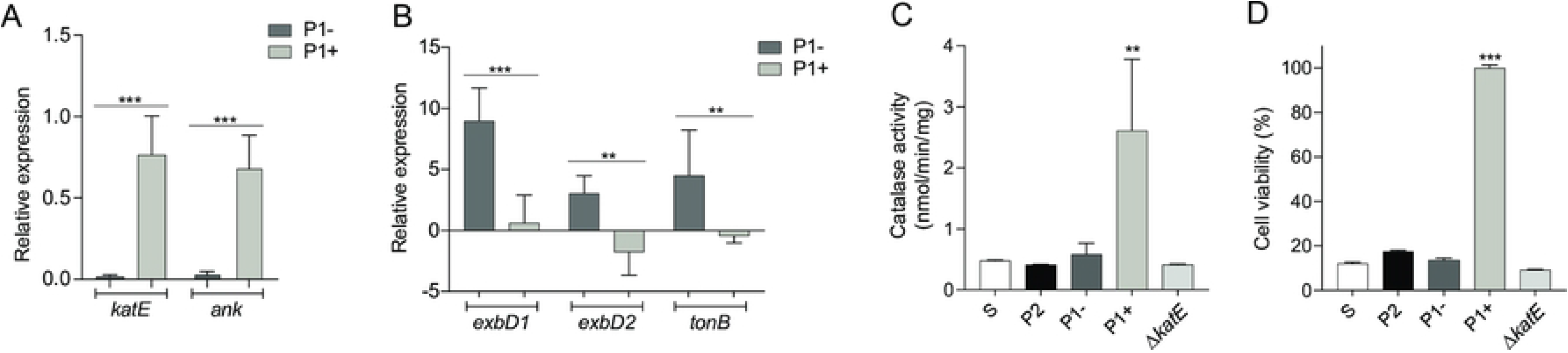
Higher resistance to oxidative stress in P1+ correlates with higher basal expression of *katE* and catalase activity. **(A-B)** Relative gene expression of *katE* and *ank* (A)*, exbD1, exbD2* and *tonB* (B) in P1+ and P1-species measured by RT-qPCR. Relative expression levels were normalized to the *flaB2* gene and compared to *L. interrogans.* **(C)** Measurement of catalase activity of *Leptospira* spp. and *Leptospira interrogans* Δ*katE* mutant. **(D)** Viability of *Leptospira* spp. and *L. interrogans ΔkatE* mutant exposed to 1 mM of H_2_O_2_ for 30 min. The percentage of *Leptospira* viability was determined by the ability of bacteria to reduce rezasurin into resorufin with or without H_2_O_2_. Unpaired two-tailed Student’s t test was used. **p<0.001, ***p<0.0001. One representative experiment (of three) is shown. Error bars represent the mean ± SD. Saprophyte (S): *L. biflexa*; P2 subclade: *L. licerasiae*, *L. fluminis*; P1-subgroup: *L. adleri*, *L. gomenensis*, *L. tipperyarensis*, *L. yasudae*; P1+ subgroup: *L. interrogans*, *L. noguchii*, *L. weilii, L. santarosai, L. mayottensis*.

In *L. interrogans*, the catalase KatE, whose expression is repressed by PerRA, detoxifies H_2_O_2_ [24]. Inactivation of *katE* resulted in an attenuation of virulence in *L. interrogans*, highlighting its role in pathogenicity [24]. We further confirmed the high basal over-expression of *katE* in P1+ by directly testing the catalase activity (Fig 4C). Interestingly, only P1+ species possess a higher basal catalase activity compared to other *Leptospira* species that contain the *katE* gene. In fact, in P1-species, the catalase activity is comparable to that of a *L. interrogans* Δ*katE* strain. To investigate the impact of a higher *katE* gene expression and catalase activity in P1+ species we analyzed the survival of P1+ and P1-species when exposed to H_2_O_2_. As expected, the survival of the P1+ species was not impaired by the presence of 1 mM of H_2_O_2,_ whereas the P1-species exhibit only 13.5% survival, a reduced survival comparable to that of the *L. interrogans* Δ*katE* mutant (Fig 4D). Interestingly, only P1+ species possess a higher basal catalase activity and better tolerance to H_2_O_2_ compared to other *Leptospira* species, even as compared with species from P and S clades. This suggests that this phenotype specifically evolved during the emergence of P1+ species. These results reveal a distinct advantage for P1+ species in resisting peroxide, despite the genetic relationship and the presence of the *katE* gene in both P1+ and P1-species.

### Genes coding for H_2_O_2_ detoxifying enzymes are less repressed by PerRA in P1+ than in P1-

We have previously shown that the catalase is repressed by PerRA in *L. interrogans* [22]. In the absence of peroxide, PerRA is under the typical metal-bound conformation prone to promoter binding and repression of *katE* gene expression [43] (Fig 5A). Similarly to a canonical PerR, PerRA releases its regulatory metal upon oxidation by H_2_O_2_, leading to DNA dissociation and alleviation of gene repression. To investigate whether PerRA-controlled gene derepression occurs in a similar manner in P1-species, we generated a Δ*perRA* mutant in *L. adleri*, used here as P1-representative species. After confirming the absence of PerRA production in this mutant by immunoblot (Fig 5B), we evaluated *katE* expression in the *L. adleri* Δ*perRA* mutant. Similarly to what is observed in *L. interrogans*, *perRA* inactivation in *L. adleri* led to the derepression of *katE* (Fig 5C) and concomitant increase in catalase activity (Fig 5D) and in tolerance to peroxide (Fig 5E). These results indicate that P1-species possess an active PerRA that represses the catalase expression and upon repression alleviation, P1-species exhibit a comparable catalase activity to that of P1+ species. We also examined whether P1-species could sense and respond to H_2_O_2_ by derepressing *katE*, as previously observed in *L. interrogans* [22]. We observed no significant up-regulation of the *katE* gene expression in P1-species in the presence of 1 mM of H_2_O_2_, while P1+ species exhibited a 3.2-fold increase in *katE* expression under these conditions (Fig 5F). However, *ahpC* and *ccp* expression was up-regulated upon exposure to H_2_O_2_ in both P1- and P1+ species but the H_2_O_2_-triggered derepression is consistently higher in P1+ species (Fig 5F). Our data show that, while PerRA can sense and respond to H_2_O_2_ by alleviating gene repression in both P1- and P1+ species, H_2_O_2_-triggered derepression is greater in P1+ than in P1-species for the same H_2_O_2_ concentration. This suggests that either PerRA_P1+_ and PerRA_P1-_ have different peroxide-sensing capacity, PerRA_P1+_ being more sensitive to oxidation by peroxide, or that PerRA_P1+_ has a lower affinity for DNA than PerRA_P1-_.

**Fig 5.**
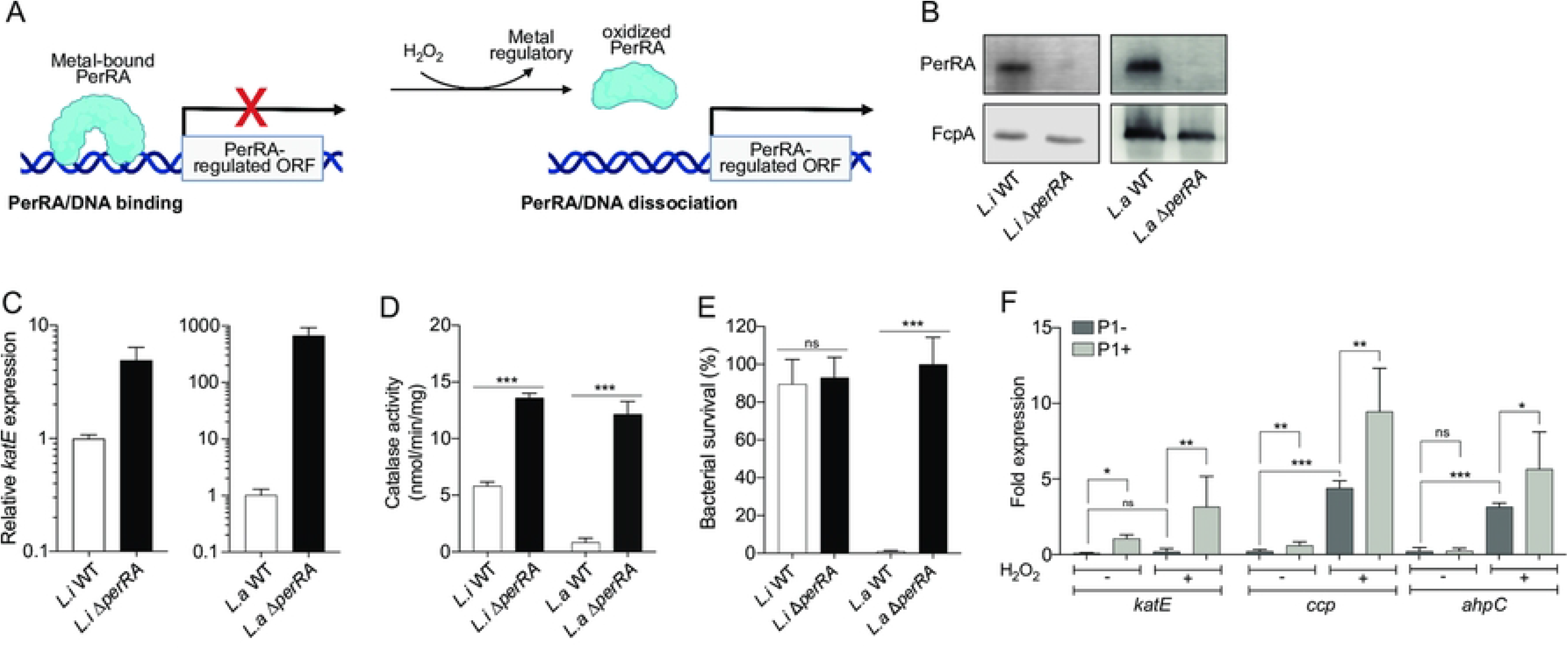
Difference of PerRA on peroxide-sensing capacity between P1+ and P1-species. **(A)** Schematic representation of the regulation of PerRA binding DNA. In absence of H_2_O_2_, PerRA is in a regulatory metal-bound conformation prone to DNA binding, resulting in peroxidase-encoding genes repression. In the presence of peroxide, PerRA is oxidized leading to the release of the regulatory metal and to a conformation switch resulting in dissociation from DNA and repression alleviation. **(B)** PerRA cellular content of the *L. interrogans* (*L.i*) WT, *L.i* Δ*perRA* mutant, *L. adleri* (*L.a*) WT and *L.a* Δ*perRA* mutant were assessed by immunoblot. FcpA is used as a control of equal loading. **(C)** Relative expression of *katE* measured by RT-qPCR in *L.i* WT*, L.i* Δ*perRA* mutant, *L.a* WT and *L.a* Δ*perRA* mutant strains upon exposure to 1 mM of H_2_O_2_ for 30 min. Relative expression levels were normalized to the *flaB2* gene and compared to *L. interrogans* without H_2_O_2_ exposure. **(D)** Measurement of catalase activity in total extracts of *L.i* WT*, L.i* Δ*perRA* mutant, *L.a* WT and *L.a* Δ*perRA* mutant strains. **(E)** Viability of *L.i* WT*, L.i* Δ*perRA* mutant, *L.a* WT and *L.a* Δ*perRA* mutant strains upon exposure to 1 mM of H_2_O_2_ for 30 min. Viability was determined by CFU and normalized with H_2_O_2_-untreated bacteria. **(F)** Relative expression of *katE*, *ccp* and *ahpC* measured by RT-qPCR in P1+ and P1-species upon exposure to 1 mM H_2_O_2_ for 30 min. Relative expression levels were normalized to the *flaB2* gene and compared to H_2_O_2_-untreated *L. interrogans*. Unpaired two-tailed Student’s t test was used. *p< 0.01, **p<0.001, ***p<0.0001, ns: non-significant. One representative experiment (of three) is shown. Error bars represent the mean ± SD. P1-subgroup: *L. adleri*, *L. gomenensis*, *L. tipperyarensis*, *L. yasudae*; P1+ subgroup: *L. interrogans*, *L. noguchii*, *L. weilii, L. santarosai, L. mayottensis*.

### Effect of a residue 89 substitution in PerRA on the H_2_O_2_-triggered derepression

In order to identify amino acid residues responsible for the weaker PerRA-mediated repression in P1+ species, we compared the protein sequences of PerRA from all P1 species (S1A Fig, Fig 6A). The DNA binding helix as well as the regulatory metal coordination site (H36, D84, H90, H92, D103), are relatively well conserved between P1+ and P1-species. We then searched for PerRA residues conserved among P1+ species but not found in all P1-species. We identified residue 89 as having a striking different distribution in P1+ and P1-species. This residue is a histidine in all P1+ species, while it is an asparagine in all P1^-^ species, (except for *L. ainazelensis* which has a proline at this position). We also observed an absence of histidine at the position 89 for the P2 and S species, which contain the *perRA* gene (except for *L. noumeaensis*). This residue is located in a loop connecting the two domains in close proximity to the regulatory metal coordination site of PerRA (Fig 6A-B), suggesting that it can participate in H_2_O_2_ sensing by PerRA. Structure predictions indicate that the presence of an Asn in position 89 does not affect the regulatory metal coordination (Fig 6B). However, prediction of the structural disordered regions in PerRA suggests that the presence of the His89 of P1+ species results in a lower probability of disordered region compared to the presence of an Asn in position 89 of PerRA_P1-_ (and other species; S1B Fig). Thus, *in silico* analysis suggests that residue 89 could have a role on PerRA differential activity.

**Fig 6.**
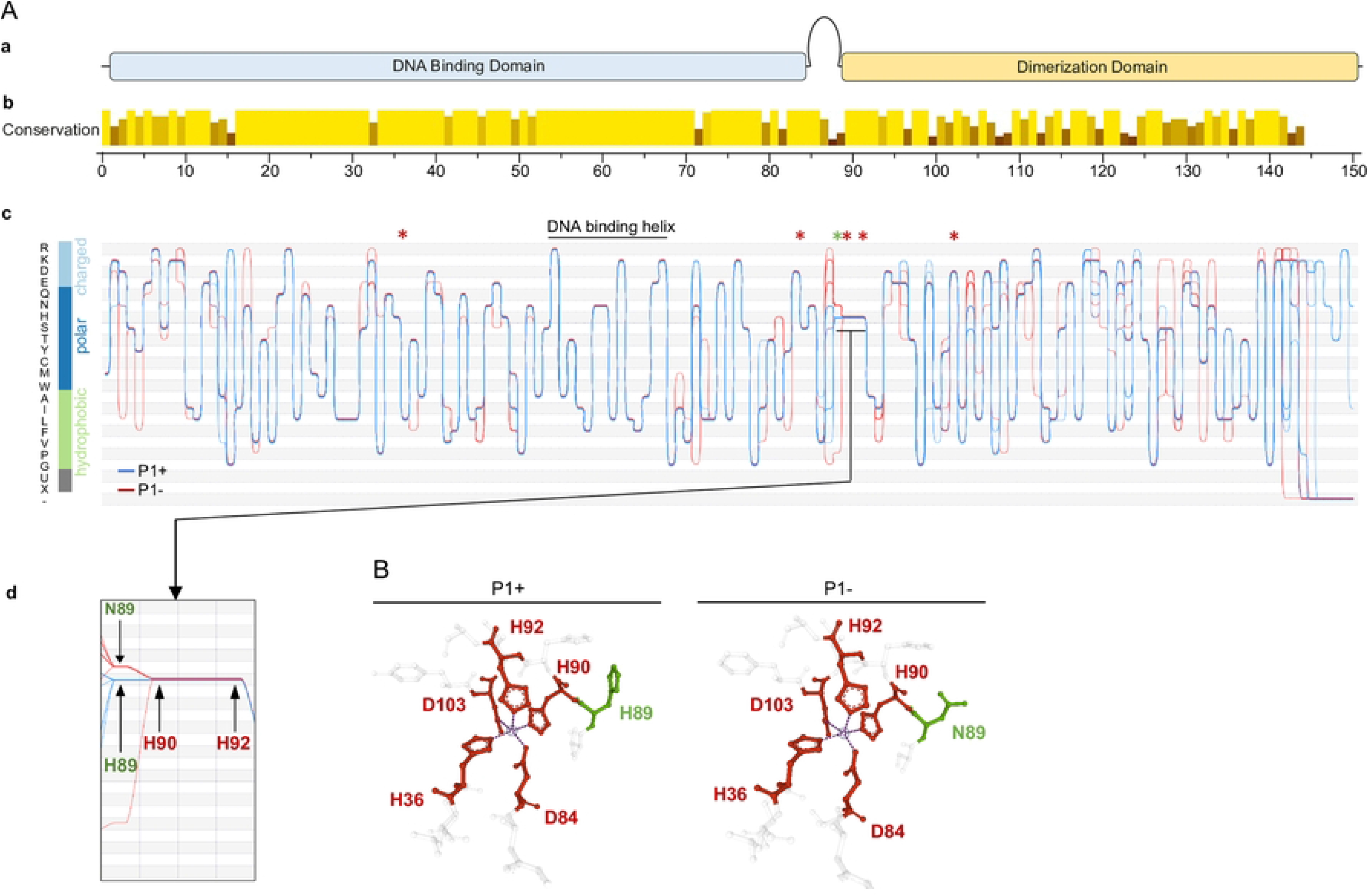
PerRA protein sequence divergence between P1+ and P1-species. **(A)** PerRA protein sequence alignment of P1+ and P1-species were performed. Schematic representation of PerRA protein sequence (a) is shown with associated domains following by the conservation of each amino acid for P1 species (b). Sequence bundles visualization of PerRA for P1+ and P1-species using Alvis software (c). Bundles plots show P1+ and P1-sequence groups in blue and red lines, respectively, represented against a Y-axis classified by amino acid hydrophobicity. Each line corresponds to one species. The curved trajectories of lines expose the conservation of residuals by converging to identical positions. Amino acids involved in regulatory metal coordination are indicated by a red asterisk and the amino acid in position 89 is indicated by a green asterisk. PerRA protein sequence alignment between the position 89 to 92 is zoomed in (d). **(B)** Detailed view of the regulatory metal binding site of PerRA of P1+ species (left) and P1-species (right). The coordination residues (H36, D84, H90, H92, and D103) are labeled in red and the metal in the regulatory metal-binding sites are represented by a sphere. Ligand coordination is symbolized by purple dashed lines. The amino acid position 89 of PerRA is labeled in green.

To further investigate the role of residue 89 of PerRA, we complemented *L. interrogans* Δ*perRA* with the heterospecific *perRA*_P1-_, with the native *perRA* from *L. interrogans* (*perRA*_P1+_) as well as with the one containing the mutation H89N (*perRA*_P1+-H89N_). We first verified that all complemented strains restored PerRA production to a similar level as *L. interrogans* WT (Fig 7A). Complementing the *L. interrogans* Δ*perRA* mutant with *perRA*_P1+,_ led to a lower *katE* repression than with *perRA*_P1+-H89N_ or *perRA*_P1-_ (Fig 7B). Interestingly, not only *katE* was dramatically more repressed by *perRA*_P1-_ but repression was not alleviated in the presence of 1 mM of H_2_O_2_. Complementing the *L. interrogans* Δ*perRA* mutant with *perRA*_P1-_ resulted in 8-fold reduction of the catalase activity and 70-fold greater survival in the presence of H_2_O_2_ than when complementing the *L. interrogans* Δ*perRA* mutant with *perRA*_P1+_ (Fig 7C-D). Therefore, differences in *katE* expression correlated with the catalase activity, in the presence and absence of 1 mM of H_2_O_2_ and with the ability of the strains to tolerate H_2_O_2_. Overall, our findings demonstrate that PerRA_P1+_ represses *katE* to a lesser extent than PerRA_P1+-H89N_ or PerRA_P1-_. It is important to note that complementation with *perRA*_P1-_ consistently led to a more dramatic effect than when complementing with *perRA*_P1+-H89N_, indicating that permutation of the residue 89 in PerRA_P1+_ could not solely recapitulate the extent of *katE* repression exerted by PerRA_P1-_. Although we observed a decrease in peroxide stress survival for *L. interrogans* Δ*perRA* expressing *perRA*_P1+-H89N_ or *perRA*_P1-_, these strains did not show attenuation of virulence in hamsters (Fig 7E).

**Fig 7.**
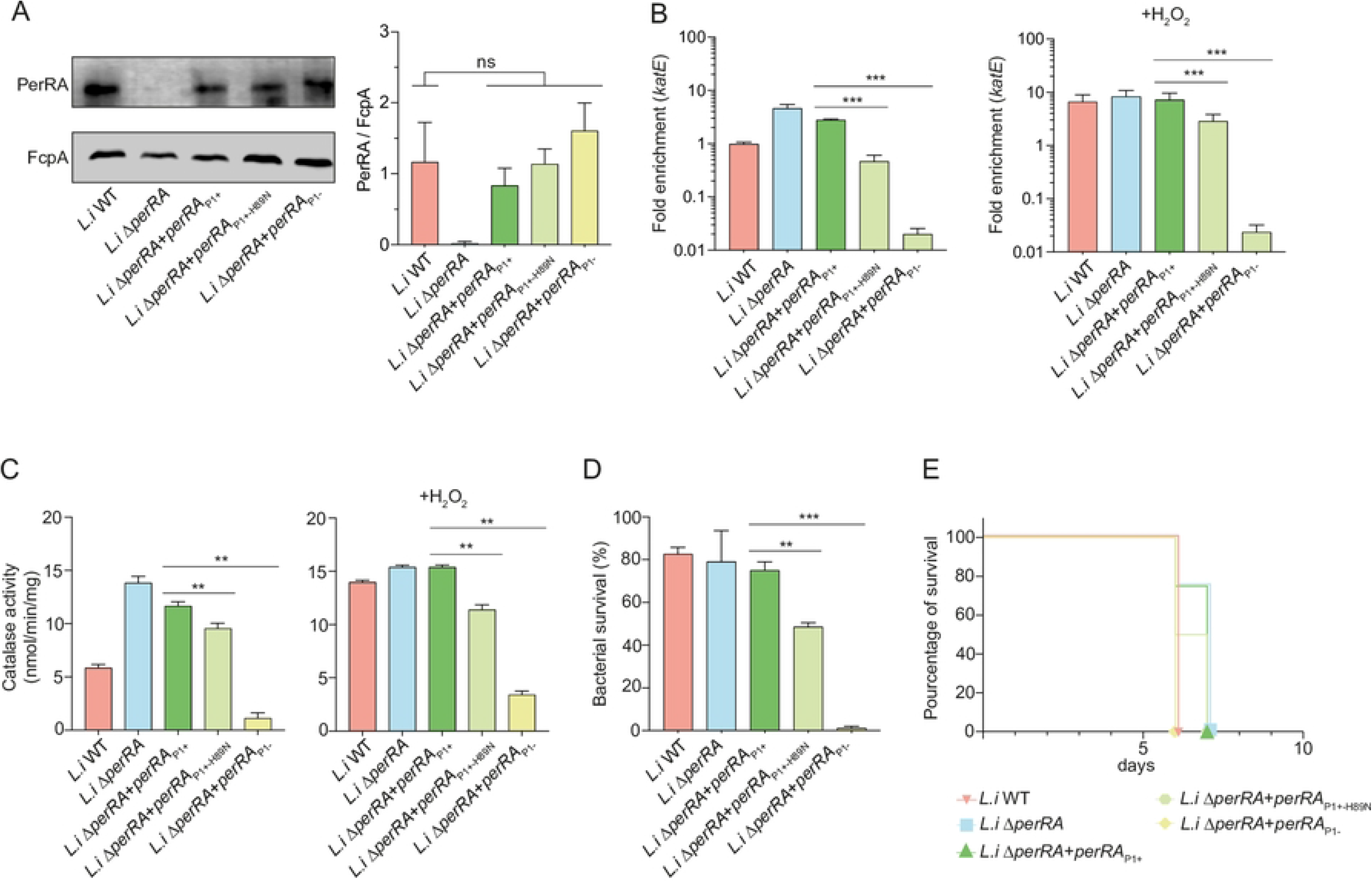
PerRA from P1+ species is associated with increased peroxide stress tolerance. **(A)** PerRA cellular content of the *Leptospira interrogans* (*L.i*) WT and Δ*perRA* mutant complemented with the indicated plasmids was assessed by immunoblot using FcpA as a control of equal loading (left panel). Densitometric quantification of PerRA was performed and normalized by FcpA amount (right panel) in *L.i* constructs. *L.i* WT and *L.i* Δ*perRA* contain the empty pMaORI-expressing vector; *L.i* Δ*perRA*+*perRA*P1+ contains the pMaORI vector bearing the *perRA* ORF of *L.i* ; *L.i* Δ*perRA*+*perRA*P1+-H89N contains the pMaORI vector bearing the *perRA* ORF of *L.i* with the single mutation H89N; *L.i* Δ*perRA*+*perRA*P1-contains the pMaORI vector bearing the *perRA* ORF of L. adleri. Data are the means and standard deviation of two independent biological replicates. **(B)** Relative expression of *katE* measured by RT-qPCR in the different *L. interrogans* (*L.i*) strains in the absence (left panel) or presence (right panel) of 1 mM of H_2_O_2_ (30 min. exposure). Relative expression levels were normalized to the *flaB2* gene and compared to *L. interrogans* WT (*L.i*). **(C)** Measurement of catalase activity in total extracts of the different *L. interrogans* (*L.i*) strains in absence (left panel) or in presence (right panel) of 1 mM of H_2_O_2_ (30 min. exposure). **(D)** Survival of the different *L. interrogans* (*L.i)* strains upon exposure to 1 mM of H_2_O_2_ for 30 min. The number of bacteria was enumerated by CFU and compared to H_2_O_2_-untreated *Leptospira*. **(E)** The virulence of the different *L. interrogans* (*L.i*) strains was assessed by infecting hamsters (n = 4) by peritoneal route with 10^6^ leptospires for each construct. For experiments in A-D, one representative experiment of three independent biological replicates is shown. Error bars represent the mean ± SD. Unpaired two-tailed Student’s t test was used. **p<0.001, ***p<0.0001, ns: non-significant.

To further assess the importance of PerRA and the H89 residue in mediating a lesser *katE* repression in P1+ species, we also complemented the *L. adleri* Δ*perRA* mutant with *L. interrogans perRA* (*perRA*_P1+_), *L. interrogans perRA* with mutation H89N (*perRA*_P1+-H89N_) or *perRA* from *L. adleri* (*perRA*_P1-_; Fig 8A). Consistent with results obtained in *L. interrogans*, complementing the *L. adleri* Δ*perRA* with *perRA*_P1+_ resulted in a lower repression of *katE* than when this mutant was complemented with *perRA*_P1-_ or with *perRA*_P1+-H89N_ (Fig 8B). Furthermore, the reduced repression of *katE* observed in the mutant *L. adleri* Δ*perRA* complemented with *perRA*_P1+_ correlated with higher catalase activity in the presence of H_2_O_2_ and an increase in tolerance to peroxide (Fig 8C-D). Since P1-species, including *L. adleri*, are rapidly cleared from hamsters, we could solely assess *Leptospira* burden in blood, kidney and liver 1-day post-infection but no significant difference was observed between the different *L. adleri* strains (Fig 8E). This indicates that the difference of catalase activity between P1+ and P1-species is not the only determinant of their difference in this model of acute infection.

**Fig 8.**
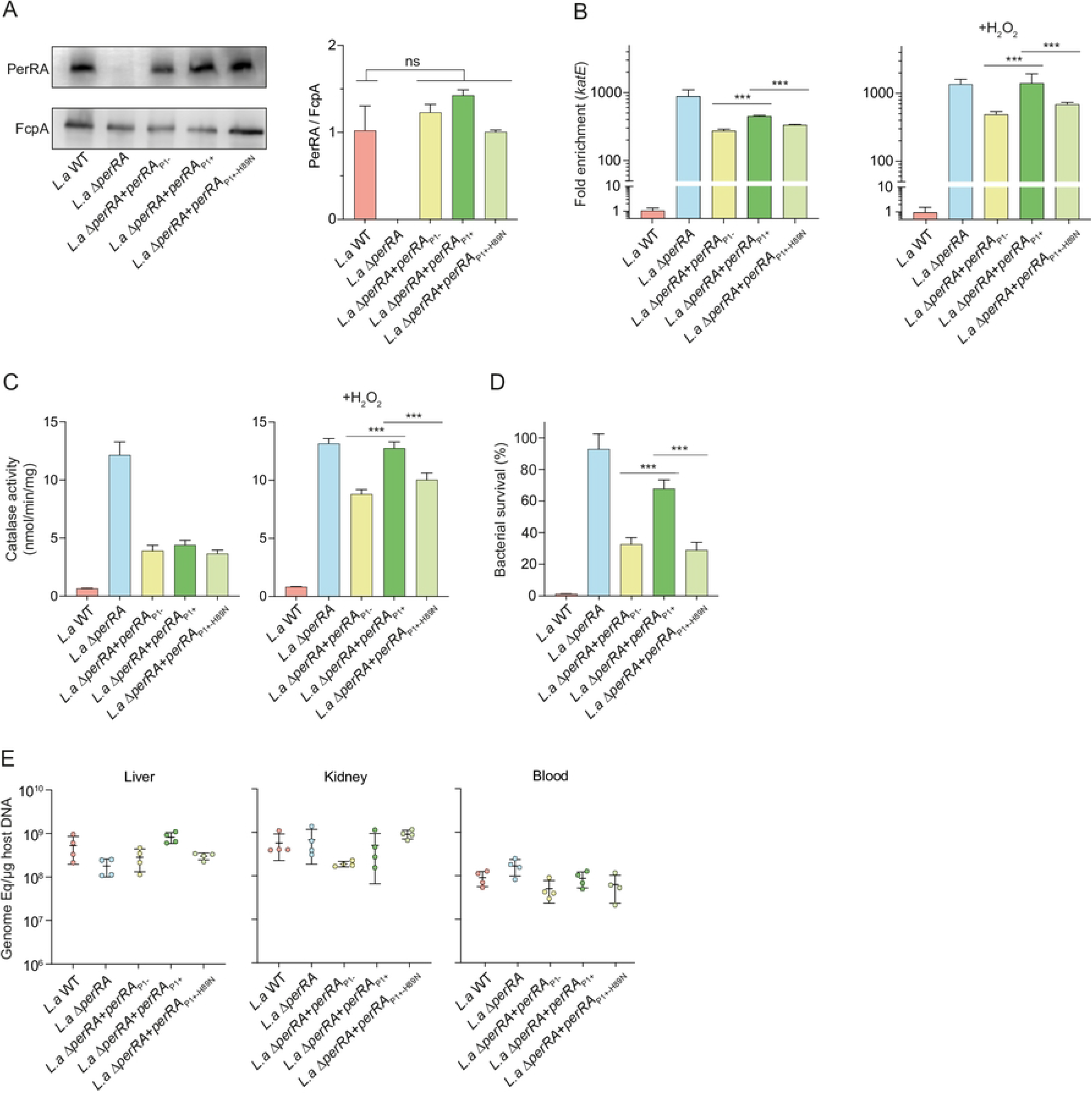
PerRA from P1+ species improve the tolerance of peroxide stress in P1-species. **(A)** PerRA cellular content of the *Leptospira adleri* (*L.a*) WT and Δ*perRA* mutant complemented with the indicated plasmids was assessed by immunoblot using FcpA as a control of equal loading (left panel). Densitometric quantification of PerRA was performed and normalized by FcpA amount (right panel) in *L.a* constructs. *L.a* WT and *L.a* Δ*perRA* contain the empty pMaORI-expressing vector; *L.a* Δ*perRA*+*perRA*_P1+_ contains the pMaORI vector bearing the *perRA* ORF of *L.a* ; *L.a* Δ*perRA*+*perRA*_P1+-H89N_ contains the pMaORI vector bearing the *perRA* ORF of *L.a* with the single mutation H89N; *L.a* Δ*perRA*+*perRA*_P1-_ contains the pMaORI vector bearing the *perRA* ORF of *L. adleri*. Data are the means and standard deviation of two independent biological replicates. **(B)** Relative expression of *katE* measured by RT-qPCR in the different *L. adleri* (*L.a*) strains in the absence (left panel) or presence (right panel) of 1 mM of H_2_O_2_ (30 min. exposure). Relative expression levels were normalized to the *flaB2* gene and compared to *L. adleri* WT (*L.a*). **(C)** Measurement of catalase activity in total extracts of *Leptospira* the different *L. adleri* (*L.a*) strains in absence (left panel) or in presence (right panel) of 1 mM of H_2_O_2_ (30 min. exposure).). **(D)** Survival of the different *L. adleri* (*L.a)* strains upon exposure to 1 mM of H_2_O_2_ for 30 min. The number of bacteria was enumerated by CFU and compared to without H_2_O_2_-untreated *Leptospira*. **(E)** The virulence of the different *L. adleri* (*L.a*) strains was assessed by infecting hamsters (n = 4) by peritoneal route with 10^8^ leptospires for each construct. After 1 day of infection, leptospiral load of hamsters infected was assessed by quantitative PCR. For experiments in A-D, one representative experiment of three independent biological replicates is shown. Error bars represent the mean ± SD. Unpaired two-tailed Student’s t test was used. ***p<0.0001, ns: non-significant.

Altogether, these findings demonstrate that difference in *katE* repression and ability to tolerate H_2_O_2_ between the P1 subgroups is mediated only by PerRA and that the residue 89 participates in tuning this repression.

## Discussion

Technological advances in sequencing have allowed the scientific community to capture genomic, epigenomic, proteomic and structural variations in species [52]. Differential gene expression studies from multiple species are one way of conducting simultaneous analysis across species. However, including multiple species requires significant processes for normalization, bias reduction and enrichment. Analysis of gene expression in multiple species is highly dependent on the evolutionary relationship between the orthologous and co-orthologous genes present, as well as a robust phylogeny. Cross-species analysis of distantly related species is complex due to the identification of homologous genes, and it may result in false predictions, grouping non-related genes into the same clusters [53]. In this study, we developed a stand-alone GUI pipeline to analyze intra-genus species using homology. The pipeline is particularly useful for understanding subtle variations in the transcriptomic repertoire of phylogenetically related species within the same genus, which may exhibit varied phenotype such as morphological features or pathogenicity. Although our pipeline is a strong tool for analysis, it has certain limitations that could be refined in the future. Orthology may be incorrectly assigned in poorly annotated or divergent genomes, which could lead to bias in the analysis. To address this, we use standardized annotation and employ two methods for homologues assignment. In addition, even though they concern only few genes from the core genome, duplications of genes in specific species also represent a significant challenge for analysis. Aside from this, we demonstrated that this unique tool is able to pinpoint some major changes in core genes expression between two groups of species. Using similar methodology, we have recently detected differential gene expression patterns between <u>Mu</u>lticellular <u>L</u>ongitudinally <u>Di</u>viding (MuLDi; *n* = 5) and rod-shaped (*n* = 5) *Neisseriaceae* cultured in the same conditions. Our analysis revealed that significantly differentially regulated genes are part of the *dcw* cluster and that this difference could be explained by the loss of the MraZ *dcw* cluster regulator in MuLDi lineage [54]. Similarly, herein in another phylum, we demonstrate and verify that two loci were differentially expressed between two groups of *Leptospira*: P1+ (highly virulent pathogens in animal models and associated with severe infections in humans) and the P1-(low virulence), that separated after the last node of evolution [19, 20]. The *katE* and *tonB* loci both belong to the PerRA regulon, and are repressed and activated, respectively, by PerRA [21, 22]. Here, we have demonstrated that the *katE* and *tonB* loci are significantly less repressed and expressed, respectively, in P1+ species than in P1-species. These two examples, clearly show the strategic fit of the methodology developed.

The methodology to study the remodelling of gene regulation during bacterial adaptation to new ecosystem or evolution of complex phenotypes is an important topic to develop. Herein, we show that there is a transcriptomic change associated with the emergence of P1+ species. The functional analysis of COG indicates a significant presence of genes encoding hypothetical proteins in both overexpressed and underexpressed genes in P1+ compared to P1-. As previously mentioned, the overexpressed genes in P1+ show a strong representation of those related to motility, a key factor in pathogenicity, as well as genes associated with signal transduction, among others. Conversely, the underexpressed genes in P1+ include those involved in metabolism and transport of amino acids, carbohydrates, lipids, and genes related to energy production. Previous studies have shown a prevalence of genes encoding energy metabolism and transport-related pathways in free-living leptospires [55]. Given that P1- are predominantly isolated from environmental samples, the underexpression of these pathways in P1+ may reflect specific adaptations associated with the transition from a primarily environmental diverse lifestyle to a more restricted host-associated one.

The detection of transcriptomic changes needs to be accompanied by studies that test and explain the mechanisms of adaptation. Previously, we have explained transcriptomic changes by the loss of a regulator [54]. Here, we were able to demonstrate the major impact of subtle adjustments in the protein sequence of the regulator PerRA. A single amino acid residue change in PerRA has a major impact on *katE* repression, leading to a higher basal *katE* expression, catalase activity and tolerance to H_2_O_2_ in P1+ species. The residue 89 is located within a loop at the hinge of the two domains in PerRA, in the vicinity of the regulatory metal coordination site (H36, D84, H90, H92, D103). The nature of this residue could impact the flexibility of this loop and thereby influence the dynamism of the metal and peroxide-triggered allosteric conformational switch. In that case, the presence of a histidine at position 89 in PerRA_P1+_ would result in a higher amount of PerRA in the metal-free conformation with low affinity for promoter region. By analogy with PerR from *Bacillus subtilis* [56], H36 and H90 are the two PerRA residues whose the oxidation by H_2_O_2_ leads to the release of the regulatory metal and dissociation from DNA. The residue at position 89 is at close proximity of the oxidized residues, therefore, it can also be speculated that the histidine 89 in PerRA_P1+_ is also a site of oxidation and, even if it does not directly participate in the regulatory metal binding, its oxidation could result in destabilizing the metal coordination. Asparagine side chain is less prone to oxidation than that of histidine, therefore the presence of an asparagine at the corresponding position in PerRA_P1-_ would lower the chance of oxidation. Other residues of PerRA might also participate in the metal and peroxide-triggered conformational switch, thereby fine-tuning gene expression. In fact, heterologous complementation experiments performed in this study showed that *katE* repression was greater when expressing *perRA*_P1-_ than when expressing *perRA*_P1+-H89N_ in the *L. interrogans* Δ*perRA* mutant. Therefore, although important, the H89N alone could not explain the higher gene repression by PerRA in P1-species. Additional divergence in PerRA protein sequence between P1+ and P1-species can be identified, including at positions 88 and 105, (Fig 6 and S1 Fig). PerRAs from P1-species possess charged amino acids at the position 88 and 105 while those of P1+ species possess polar or hydrophobic amino acids. We can infer that such differences might also participate in the difference of PerRA activity between P1+ and P1-species. Thorough biochemical characterization of the PerRAs from the two P1 subgroups will be necessary to understand the exact role of these different residues and to verify the hypotheses mentioned above.

Catalase has been shown to be required for *Leptospira* virulence [24]. *KatE* gene is present in all species from the P1 subclade but this study suggests that a reduced repression of *katE* resulting in a higher catalase activity may have emerged as an evolutionary advantage for highly virulent species. Together with a higher catalase activity, P1 + species have a reduced expression of a cluster encoding the TonB-dependent energy transduction system. TonB-dependent transporters are often involved in the uptake of ferric siderophores, among other molecules. Accumulation of iron upon peroxide stress worsens oxidative damage because it favors the production of hydroxyl radicals through the Fenton reaction. Lowering iron uptake together with increasing peroxide breakdown by catalase would allow P1+ species to be better equipped to resist the oxidative stress encountered when infecting a host. Using animal model of acute infection, we were not able to show a clear reduction of virulence of P1+ species or an increase in the ability of P1-species to colonize a mammalian host when *katE* repression was modulated by heterologous complementation. Core gene expression comparison across P1 subclade have identified additional genes that are overexpressed in P1+ species, and some of the factors encoded by these genes might be determinant for *Leptospira* virulence in addition to the catalase. In addition, fine-tuning of gene expression control by PerRA may be important at the scale of lineage evolution but too subtle to reproduce in animal models. In addition, it can be implicated in other part of the life cycle of P1+ species important for their maintenance in the hosts population (chronic colonization of asymptomatic reservoir) not tested here.

In summary, this study revealed a method and a user-friendly GUI to study intra-genus core-gene expression change and exemplify it by studying a regulatory network in pathogenic *Leptospira* species. We demonstrate that we can detect efficiently true transcriptomic changes by showing that a common regulator, PerRA, has different properties between P1+ and P1-species. The reduced PerRA-mediated repression observed in P1+ species facilitates their survival against oxidative stress, probably enabling them to resist harmful oxidants produced by the host’s innate immune response and promoting a permissive host infection. This example highlights the importance of combining genomic comparisons with gene expression profiling to fully understand the emergence of complexes phenotypes at the genus scale.

## Acknowledgements

We thank Farah Martin for help with plasmid constructs, Robert Gaultney for RNA extraction. This research was supported by the Canadian Institutes of Health Research (CIHR) (#450862) (FJV); by the Institut Pasteur through grant PTR 30-2017 (MP and FJV) and by Institut Pasteur & INRS-Centre Armand-Frappier through their Pasteur International Joint Research Units program “LEptospirosis Pasteur NETwork (LePNet)” (MP and FJV). CN received a Ph.D. studentship Calmette & Yersin from the Institut Pasteur International Network. AGG was funded by a PTR2019-310 grant (NB). FJV received a Junior 1 and Junior 2 research scholar salary award from the Fonds de Recherche du Québec—Santé. The funders had no role in study design, data collection and analysis, decision to publish, or preparation of the article.

---

**S1 Figure. PerRA protein sequence in *Leptospira spp*. (A)** Alignment of PerRA protein sequence in *Leptospira* spp. and (**B)** Disorder tendency score of each residue in the PerRA protein are shown for P1+ and P1-species using IUPred2A. Higher values correspond to higher probabilities of disorder. The residue at the position 89 is represented by a green asterisk.

**S1 Table.** The table lists *Leptospira* species used in this study.

**S2 Table.** The table lists all primers used in this study.

**S3 Table.** The table lists of plasmids used in this study.

**S4 Table.** Excel tables presenting the GeneAssociation tables obtained using GH (sheet 1) or NX (sheet 2), the CountComparisonOutput obtained using featurescounts for GH (sheet 3) or NX annotations (sheet 4). Sheet 5 presents all the DeSeq2 statistical analyses results (including the sequence and name of proteins).

**S5 Table.** Excel tables presenting the data relative to the COG analysis performed through eggNOG mapper. Considering the default values of an e-value of 0.001 and confidence score of 60, 122 out of 202 significantly differentially downregulated genes in P1+ were successfully annotated (sheet 1). Similarly, for significantly upregulated genes in P1+, 160 out of 319 were assigned a functional annotation (sheet 2). Please note that few proteins may have multiple functional annotations, and these were considered in the counting to express representativeness of each category (sheet 3).

**S6 Table.** Table presenting regulated-genes in PerRA regulon in P1+ species compared to P1-species.

## Notes

### Competing Interest Statement

The authors have declared no competing interest.

